# Mitofusin 2 regulates neutrophil adhesive migration and the actin cytoskeleton

**DOI:** 10.1101/608091

**Authors:** Wenqing Zhou, Alan Y. Hsu, Yueyang Wang, Tianqi Wang, Jacob Jeffries, Xu Wang, Haroon Mohammad, Mohamed N. Seleem, David Umulis, Qing Deng

## Abstract

Neutrophils rely on glycolysis for energy production. How mitochondria regulate neutrophil function is not fully understood. Here, we report that mitochondrial outer membrane protein Mitofusin 2 (Mfn2) regulates neutrophil homeostasis in vivo. *Mfn2*-deficient neutrophils are released from the hematopoietic tissue and trapped in the vasculature in zebrafish embryos. Human neutrophil-like cells deficient with MFN2 fail to arrest on activated endothelium under sheer stress or perform chemotaxis. Deletion of Mfn2 results in a significant reduction of neutrophil infiltration to the inflamed peritoneal cavity in mice. Mfn2, but not Mfn1, -null mouse embryonic fibroblast cells have altered actin structure and are impaired in wound closure. MFN2-deficient neutrophil-like cells display heightened intracellular calcium levels and Rac activation after chemokine stimulation. Mechanistically, MFN2 maintains mitochondria-ER interaction. Restoring mitochondria-ER tether rescues the chemotaxis defect and Rac activation resulted from MFN2 depletion. Finally, inhibition of Rac restores chemotaxis in MFN2-deficient neutrophils. Altogether, we identified that MFN2 regulates neutrophil migration via suppressing Rac activation and uncovered a previously unrecognized role of MFN2 in regulating the actin cytoskeleton.

## Introduction

Neutrophils, the most abundant circulating leukocytes in humans, constitute the first line of host defense. Upon stimulation by either pathogen or host-derived proinflammatory mediators, neutrophils are recruited to inflamed tissue using spatially and temporally dynamic intracellular signaling pathways. Activation of the surface receptors, primarily the G-protein-coupled receptors (de Oliveira et al., 2016; Gambardella and Vermeren, 2013; Mocsai et al., 2015; Pantarelli and Welch, 2018), leads to the activation of phosphatidylinositol 3-kinases (PI3K) that produces phosphatidylinositol (3,4,5)P3 and activates small GTPases such as Rac. Rac promotes actin polymerization at the leading edge and drives cell migration (Futosi et al., 2013). In parallel, G-protein-coupled receptors activate phospholipase C, which generates IP3 and promotes the Ca^2+^ release from intracellular stores (Tsai et al., 2015). Although intracellular Ca^2+^ is a well characterized second messenger that activates Rac and regulates cell migration in slowly migrating cells (Price et al., 2003), its role in neutrophil migration is less clear.

Cell migration requires the coordination of multiple cellular organelles, including mitochondria. Mitochondria carry out oxidative phosphorylation to produce ATP, regulate the intracellular redox status and distribution of Ca^2+^ that can regulate cell migration. In addition, mitochondria morphology changes via fusion and fission (Campello and Scorrano, 2010) to adapt to changing metabolic needs under different conditions. Mitochondria fission promotes cell migration by providing mitochondria and ATP at energy demanding sites such as the protrusion or the uropod (Campello et al., 2006; Zhao et al., 2013).

In neutrophils, mitochondrial biology is distinct and contradictory. The Warburg effect is documented in neutrophils that they primarily use glycolysis for ATP generation (Borregaard and Herlin, 1982). Neutrophils have a relative low number of mitochondria, low respiration rates and low enzymatic activity of the electron transport chain (Peachman et al., 2001).

However, disrupting mitochondrial membrane potential by pharmacological inhibitors abolished chemotaxis of primary human neutrophils (Bao et al., 2015; Bao et al., 2014; Fossati et al., 2003). Although mitochondria-derived ATP possibly regulates neutrophil chemotaxis in vitro (Bao et al., 2015), removal of extracellular ATP improves neutrophil chemotaxis in vivo (Li et al., 2016). These conflicting reports prompted us to search for mechanisms delineating the role of mitochondria in neutrophil migration outside the realm of ATP or cellular energy (Bi et al., 2014; Schuler et al., 2017; Zanotelli et al., 2018).

Human neutrophils are terminally differentiated and undergo apoptosis within 24 hours in culture and thus are not genetically tractable. We have overcome this hurdle by developing a neutrophil-specific knockout platform in zebrafish (Zhou et al., 2018a). The zebrafish is a suitable model for neutrophil research because of its highly conserved innate immune system. In our previous work, we have confirmed the requirement of mitochondrial membrane potential and electron transport chain in the migration of zebrafish neutrophils (Zhou et al., 2018a). In addition, we have visualized a highly fused tubular network of mitochondria in zebrafish neutrophils, which is consistent with a previous report investigating primary human neutrophils (Maianski et al., 2002). Here we present evidence that Mitofusin 2 (Mfn2) regulates Rac activation to coordinate neutrophil adhesion and migration. In addition, we reveal a previously unknown function of Mfn2 in regulating the actin cytoskeleton, contributing to the understanding and management of patients with Mfn2-related mitochondrial diseases.

## Results

### Neutrophils depleted with *mfn2* accumulate in zebrafish vasculature

To address whether a highly fused mitochondrial network benefits neutrophil migration, we generated zebrafish transgenic lines with neutrophil specific deletion of proteins that regulate mitochondrial shape. Mitofusins, Mfn1 and Mfn2, are required for mitochondrial outer membrane fusion (Chen et al., 2003) and Opa1 regulates inner membrane fusion (Song et al., 2007). In embryos with *mfn2* deletion in neutrophils, *Tg(lyzC:Cas9-mfn2 sgRNAs)^pu23^*, a majority of neutrophils circulate in the bloodstream (Fig. 1a, b and Supplementary Movie 1). This is in sharp contrast to the control or the wild-type embryos in which over 99% of neutrophils are retained in the caudal hematopoietic tissue or in the head mesenchyme (Harvie and Huttenlocher, 2015). This abnormal distribution of neutrophils was further confirmed in a second transgenic line expressing different sgRNAs targeting *mfn2*, *Tg(lyzC:Cas9-mfn2 sgRNAs#2) ^pu24^* (Fig. 1a, b and Supplementary Movie 2). Neutrophils were sorted from both lines and their respective loci targeted by the 4 sgRNAs were deep sequenced. The overall mutation frequency ranged from 24% to 60%. In contrast, circulating neutrophils were not observed in embryos expressing sgRNAs targeting *opa1,* although the velocity of neutrophil migration in the head mesenchyme was significantly reduced (Supplementary Fig. 1 and Supplementary Movie 3), indicating that decreased neutrophil retention in tissue is not simply due to defects in mitochondrial fusion.

**Figure 1.**
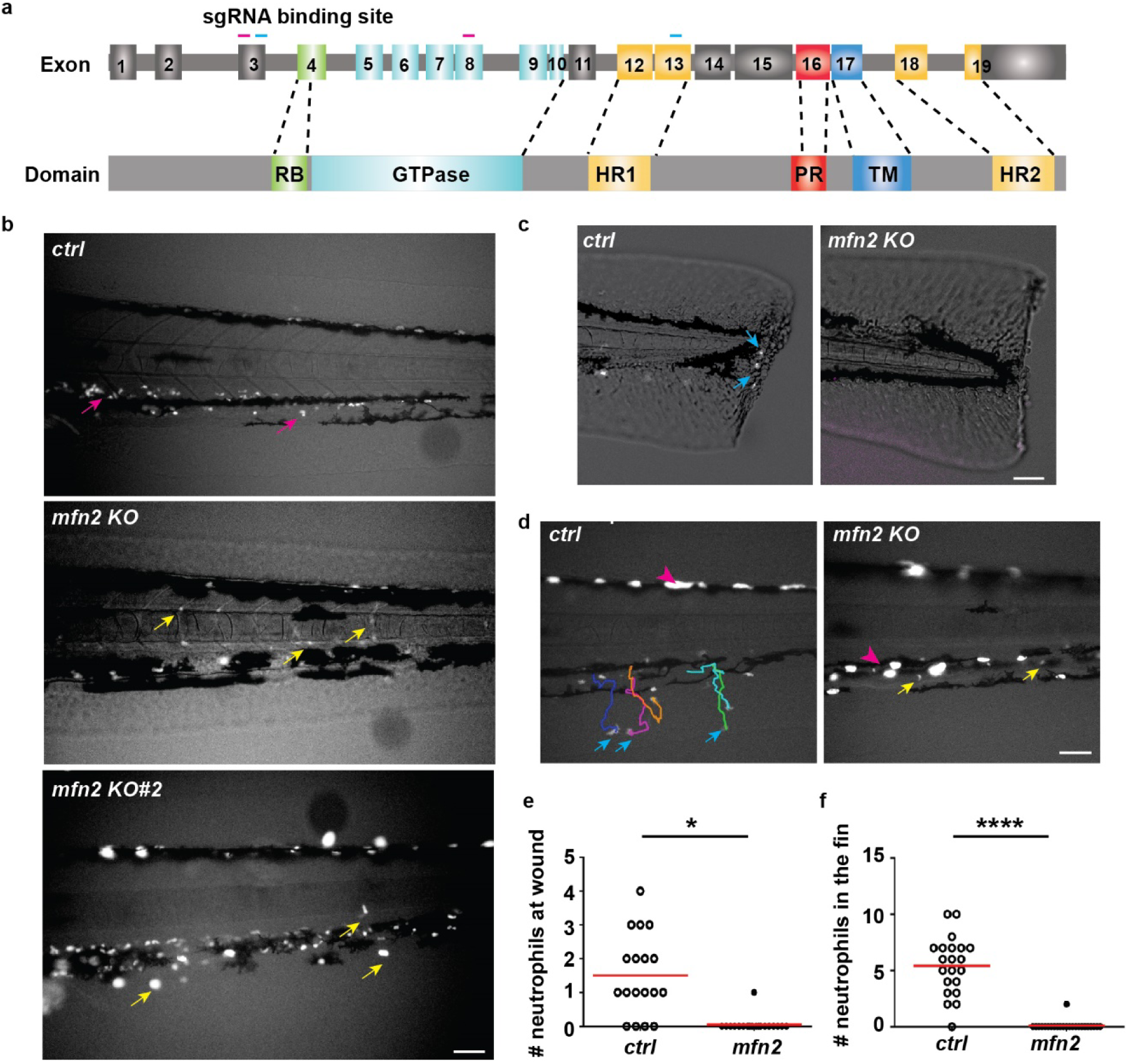
Mfn2 regulates neutrophil homeostasis in zebrafish. a) Schematics of the gene structure and protein domains of the zebrafish *mfn2* gene. The first set of sgRNAs (purple lines) targets exon 3 and exon 8 in the forward strand, and the second set (blue lines) targets exon 3 and exon 13 in the forward strand. b) Representative images of neutrophils in the zebrafish trunk of the transgenic lines with neutrophil specific *mfn*2 disruption at 3 dpf. Magenta arrows: neutrophils in the caudal hematopoietic tissue; yellow arrows: neutrophils in the vasculature. c) Representative images and e) quantification of neutrophil recruitment to the wound edge at 1 hpw. Blue arrows: neutrophils migrated to the wound. d) Representative tracks and f) quantification of neutrophil recruitment to the fin at 30 min post LTB_4_ treatment. Blue arrows: neutrophils in the fin. One representative result of three biological repeats is shown in d and f. n>20 fish embryos in each group were quantified in e and f. *, p<0.05, ****, p<0.0001, unpaired *t*-test. Scale bar: 50 µm.

Next, we determined whether neutrophils in the vasculature were able to respond to acute inflammation induced by a tail transection or perform chemotaxis to LTB_4_. Significant defects in both assays were observed in the line with neutrophil specific *mfn2* deletion (Fig. 1c-f and Supplementary Movie 4). Taken together, *mfn2* regulates neutrophil tissue retention and extravasation in zebrafish.

### MFN2 regulates adhesion and migration in of neutrophils in vitro and in vivo

To get to the mechanism how Mfn2 regulates neutrophil migration, we knocked down MFN2 in human neutrophil-like cells, HL-60, using shRNAs and obtained two individual lines with 80% and 50% reduction of MFN2 (Fig. 2a). No differences in cell viability, differentiation, surface expressions of integrins or selectin ligands were noted (Supplementary Fig. 2a-g). Both lines are significantly deficient with chemotaxis towards fMLP on collagen coated 2-dimensional surfaces (Fig. 2b, c). The defect in chemotaxis was rescued by reconstitution with a shRNA-resistant MFN2 in MFN2 knockdown cells (Fig. 2d-f and Supplementary Movie 5), supporting that the shRNA specifically targeted *MFN2*. In addition, acute reduction of MFN2 in dHL-60 resulted in similar chemotaxis defects (Fig. 2g, h and Supplementary Movie 6), suggesting that this defect is not due to nonspecific secondary effects associated with chronic MFN2 depletion. Under shear stress, a majority of *MFN2*-deficient cells failed to firmly adhere to activated endothelial cells (Fig. 2i, j and Supplementary Movie 7), recapitulating the phenotype in zebrafish where neutrophils depleted with *mfn2* failed to adhere to the vasculature. Intriguingly, knocking down MFN1, which shares a similar structure and function with MFN2, in dHL-60 cells, did not affect chemotaxis (Supplementary Fig. 3a-c and Supplementary Movie 8). To investigate whether Mfn2 is required for neutrophil infiltration in mice, we bred *Mfn2* flox/flox mice (Chen et al., 2007) with the *S100A8-Cre* strain (Abram et al., 2013) for neutrophil specific depletion. With 50% of *Mfn2* transcript reduction in neutrophils obtained in this strain, significant reduction of neutrophil infiltration into the inflamed peritoneal cavity was observed (Fig. 2 k-m). Blood cell composition was not altered by Mfn2 depletion (28% and 32% granulocytes in the *Cre*^+^ and *Cre*^-^ lines, respectively), consistent with a previous report that *Mfn2* does not regulate blood cell development under hemostatic conditions (Luchsinger et al., 2016). Therefore, MFN2 is required for neutrophil chemotaxis and infiltration in mammals.

**Figure 2.**
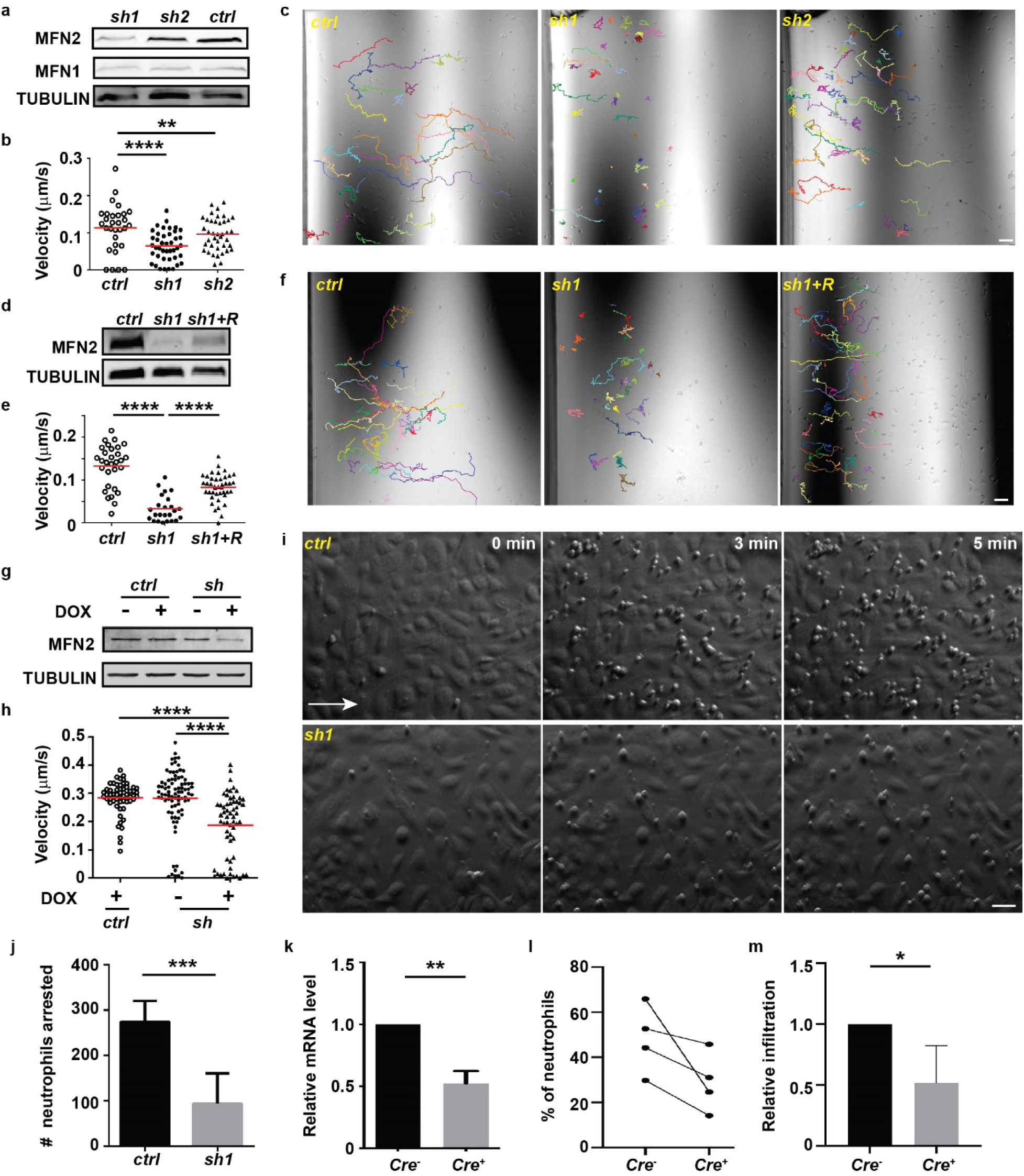
MFN2 regulates neutrophil migration in vitro and in vivo. a) Western blot determining the expression level of MFN2 in indicated cell lines. *ctrl*: standard control; *sh1*: shRNA targeting MFN2; *sh2*, another shRNA targeting MFN2. b) Quantification of velocity and c) Representative images with individual tracks of neutrophil chemotaxis to fMLP. d) Western blot showing the expression level of MFN2 in indicated cell lines. e) Quantification and f) Representative images with individual tracks of neutrophils migrating toward fMLP. g) Western blot of MFN2 in indicated cell lines with or without doxycycline induction. h) Quantification of velocity of neutrophil chemotaxis to fMLP. i) and j) Adhesion of neutrophils under sheer stress. HUVEC monolayer was activated with TNFα and neutrophils were flowed on top of the monolayer for 5 min. i) Representative images showing neutrophils arrested by HUVEC at indicated time points. White arrow: flow direction. j) Quantitation of numbers of neutrophils arrested at 5 min. k) The relative mRNA level of *Mfn2* in mice neutrophils isolated from *Mfn2^flox/flox^; S100A8:Cre+* or the control *Mfn2^flox/flox^; S100A8:Cre-*littermates. l) Percentage of neutrophil in the peritoneal cavity in indicated mice. m) Relative neutrophil infiltration to peritoneal cavity. Percentage of neutrophils in the lavage was normalized to that in sex-matched littermates in each experiment. One representative result of three (a-h) or two (i-j) biological repeats is shown. Data are pooled from three (k and j) or four (l and m) independent experiments. n>20 cells are tracked and counted in c, d, e, f, i, h. **, p<0.01; ****, p<0.0001, one-way ANOVA (b, e, h). **, p<0.01, ***, p<0.001, unpaired *t*-test (j, and k). *, p<0.05, paired *t*-test (m). Scale bar: 50 µm.

### Mfn2 regulates the actin cytoskeleton and migration of mouse embryonic fibroblasts

In addition to neutrophils, we investigated the function of Mfn2 in mouse embryonic fibroblasts (MEF). *Mfn2-null* MEFs are round with enriched actin filaments and Paxillin in the cell cortex, whereas *wt* MEFs are elongated with stress fibers when plated on ligand-coated or uncoated substrates (Fig. 3a,b and Supplementary Fig. 3d). *Mfn1*-deficient MEFs are round but retained stress fibers (Supplementary Fig. 3d-f). The significant changes in actin organization suggest that *Mfn2-null* MEFs may migrate differently. Indeed, MEFs deficient with Mfn2 migrated slower when compared to *wt* cells during wound closure (Fig. 3c, d and Supplementary Movie 9). In addition, after plating, *wt* MEFs extended transient filopodia and lamellipodia and eventually elongated, whereas *Mfn2-null* MEFs generated extensive membrane ruffles and retained the circular shape (Fig. 3e and Supplementary Movie 10). *Mfn1-null* cells spread similarly to the *wt* cells (Supplementary Fig. 3g). In summary, Mfn2 modulates the actin cytoskeleton and cell migration in MEFs.

**Figure 3.**
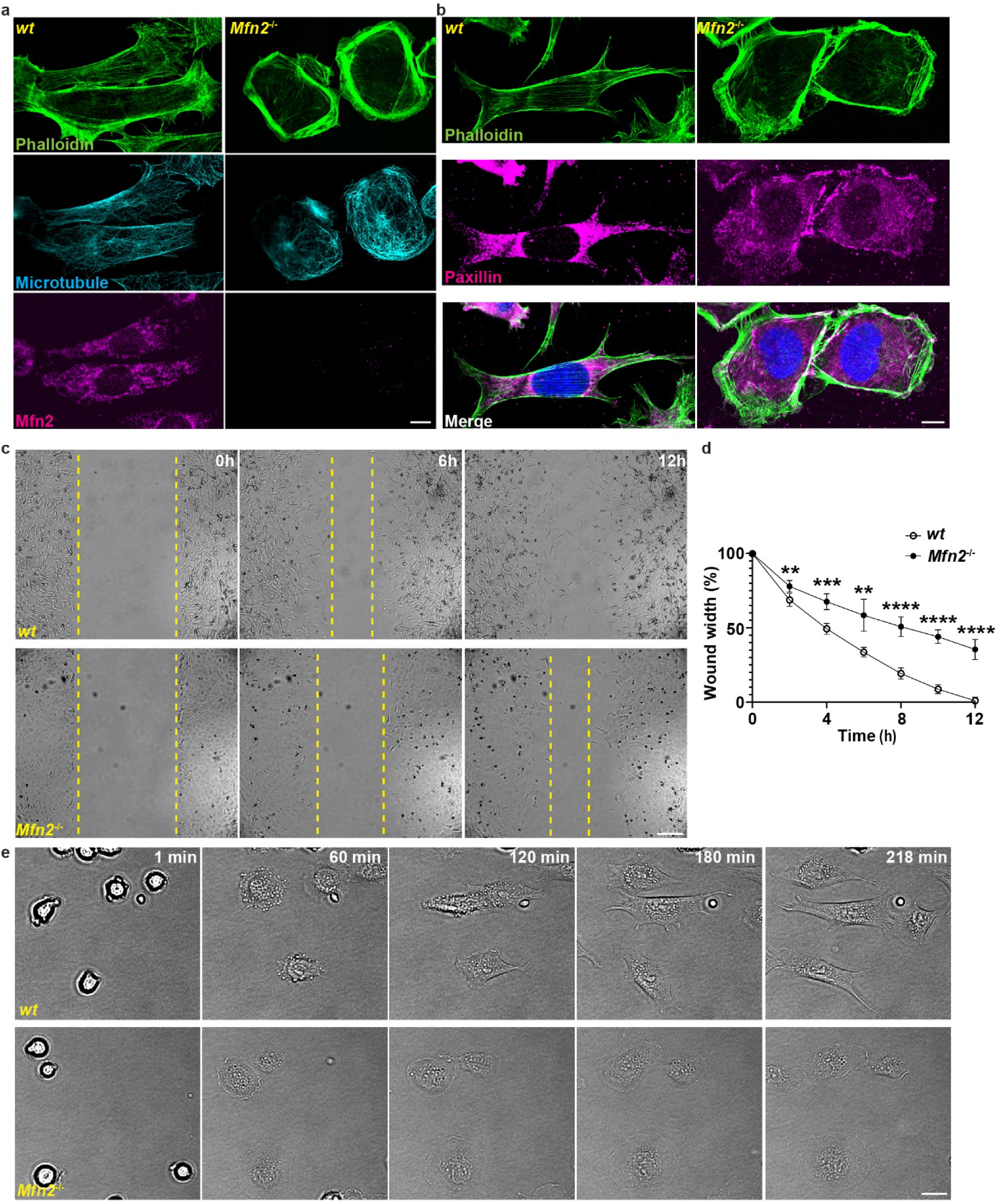
Mfn2 regulates cytoskeleton organization and cell migration in MEFs. a) Immunofluorescence of F-actin (phalloidin), microtubule and Mfn2 in *wt* and *Mfn2-null* MEFs. b) Immunofluorescence of F-actin and Paxillin in wt and *Mfn2-null* MEFs. c) Representative images of wt and *Mfn2-null* MEFs during wound closure at indicated time points. Yellow dash lines: wound edge. d) Quantification of the wound area in wt and *Mfn2-null* MEFs during wound closure. e) Representative images of wt and *Mfn2-null* MEFs during cell spreading at indicated time points. One representative result of three biological repeats is shown in a-e. **, p<0.01, ***, p<0.001, ****, p<0.0001, unpaired *t*-test. Scale bar: 10 µm (a and b), 20 µm in e and 200 µm in c.

### MFN2 regulates cell migration through maintaining mitochondria-ER tether in dHL-60 cells

We have demonstrated a role of Mfn2 in regulating cell migration in different model systems. Next we want to investigate the underlying molecular mechanisms. Mfn2 localizes to both the mitochondria and the ER membrane and regulates the tethering between the two organelles in MEF cells (Naon et al., 2016). In dHL-60 cells, MFN2 colocalized with both the mitochondria and the ER, with Manders’ colocalization coefficiencies of 0.60±0.085 and 0.69±0.13, respectively (Fig. 4a, b). Mitochondria also colocalized with the ER (Manders’ colocalization coefficiency 0.52±0.097) and distributed throughout the cell body (Fig. 4c, d). The morphology of the mitochondria and the ER was further visualized using electron microscopy in dHL-60 cells (Supplementary Fig. 4). When MFN2 was inhibited, mitochondria lost their structure and interaction with ER, and formed a cluster in the middle of the cell body (Fig. 4c, d). Interestingly, reconstitution of the MFN2 knockdown dHL-60s with an artificial tether (which bridges mitochondria and ER independent of MFN2) (de Brito and Scorrano, 2008) restored the morphology and structure of mitochondria in MFN2-deficient dHL-60 (Fig. 5a-d). Furthermore, expression of the artificial tether rescued the chemotaxis defect in MFN2-deficient dHL-60 cells (Fig. 5d-f and supplementary Movie 11). Taken together, MFN2 controls cell migration by bridging mitochondria and ER in dHL-60 cells.

**Figure 4.**
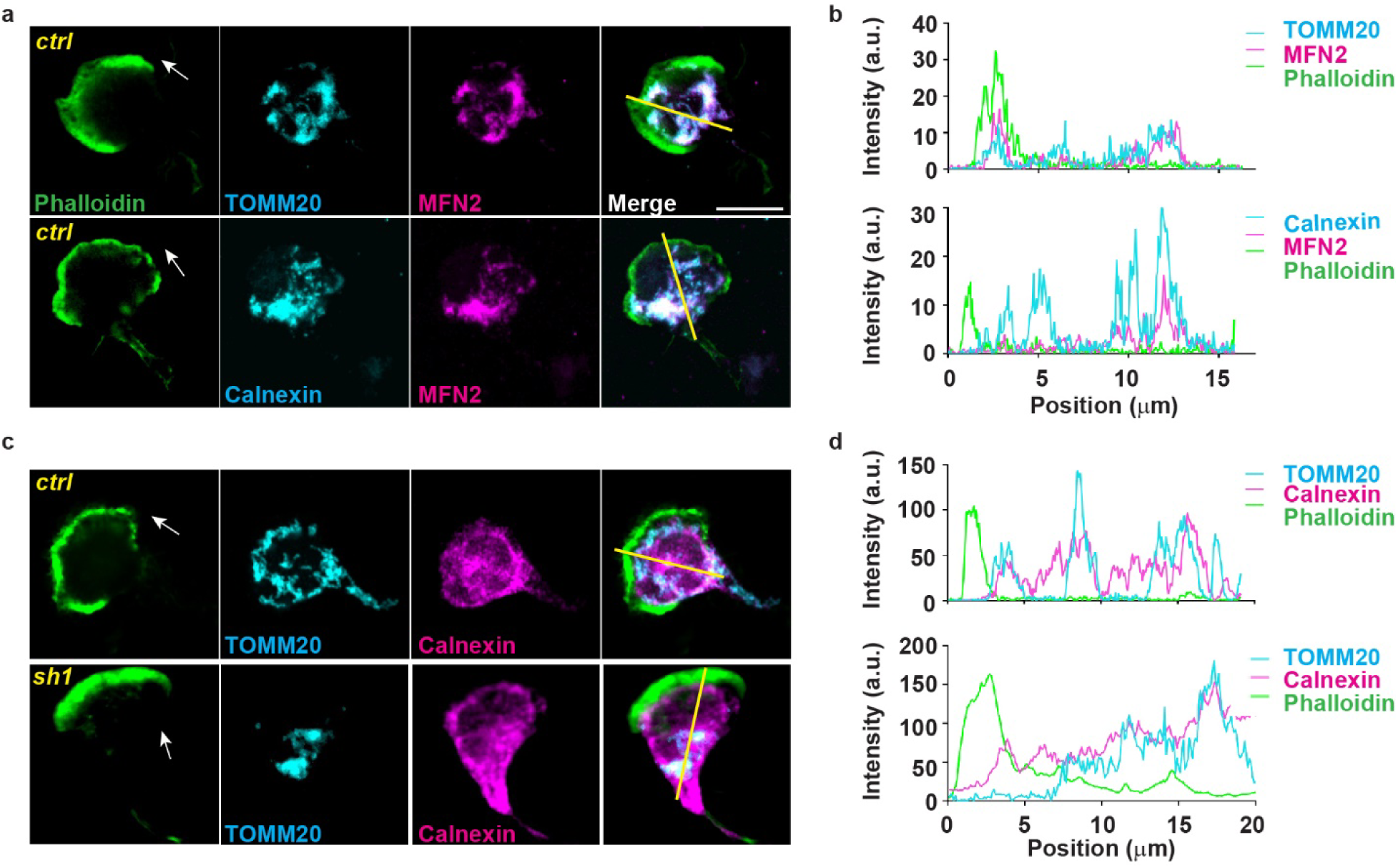
MFN2 regulates ER-mitochondria tethering. a) Immunofluorescence of, mitochondria (TOMM20), MFN2 and ER membrane (Calnexin) in indicated cells 3 min post fMLP stimulation. Cells were stained also with phalloidin to reveal F-actin. Arrows: direction of cell polarization. b) Plot profiles of the fluorescence intensity (MFI) along the corresponding yellow lines in a. c) Immunofluorescence of, mitochondria and ER membrane in indicated cells 3 min post fMLP stimulation. Arrows: direction of cell polarization. d) Plot profiles of the fluorescence intensity (MFI) along the corresponding yellow lines in c. One representative result of three biological repeats was shown in a-d. Scale bar: 10 µm.

**Figure 5.**
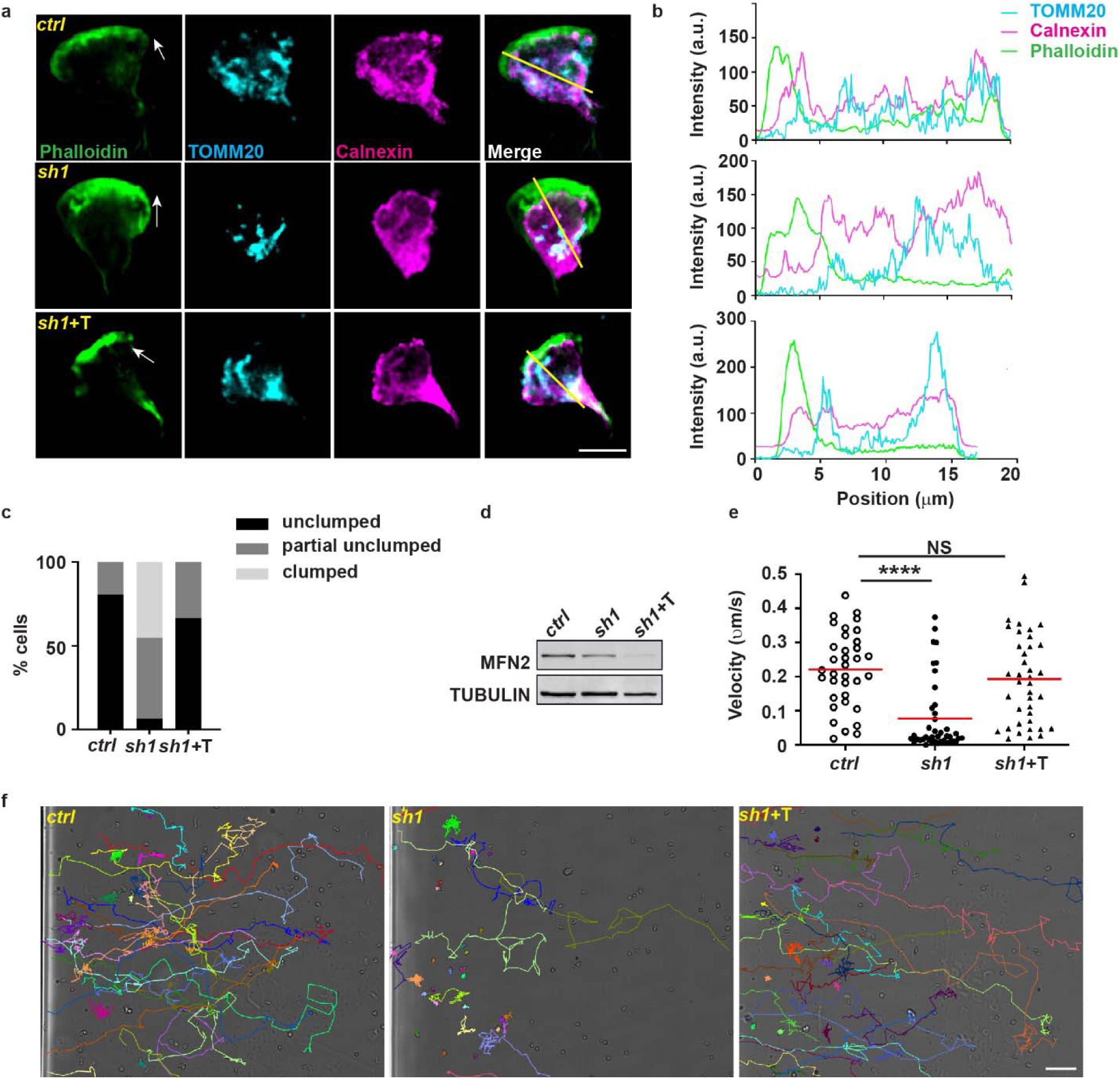
MFN2 regulates intracellular Ca^2+^ and suppresses RAC over-activation in dHL-60 cells. a) Immunofluorescence of mitochondria (TOMM20), and ER membrane (Calnexin) in indicated cells 3 min post fMLP stimulation. Arrows: direction of cell polarization. b) Plot profiles of the fluorescence intensity (MFI) along the corresponding yellow lines in a. c) Quantification of clumped mitochondria in indicated cell lines. d) Western blot of MFN2 in indicated cell lines. sh1+T: HL-60 cells with MFN2-sh1 and synthetic tether construct. d) Quantification of neutrophil velocity and e) Representative images of individual tracks of neutrophils migrating to fMLP. One representative result of three biological repeats is shown in a, b, d, e, and f. Data are pooled from three independent experiments in c. n>20 cells are quantified or tracked in c, e, and f. ****, p<0.0001, one-way ANOVA in f. Scale bar: 10 µm in a, 100 µm in e.

### MFN2 suppresses RAC activation in dHL-60 cells

The close proximity of the ER and the mitochondria regulates multiple cellular signaling pathways including calcium homeostasis (de Brito and Scorrano, 2008). Indeed, *MFN2*-deficient dHL-60 cells exhibited higher levels of Ca^2+^ in the cytosol and reduced levels in the mitochondria after fMLP stimulation (Fig. 6a, b). ATP levels were not affected in the MFN2 knockdown dHL-60 cells (Supplementary Fig. 5a), in line with the observation that mitochondria are not a major source of ATP in neutrophils (Amini et al., 2018; Borregaard and Herlin, 1982). Mitochondrial membrane potential and the ROS level in mitochondria were also reduced when MFN2 was depleted, especially after fMLP stimulation (Supplementary Fig. 5b-e). This may due to the altered level of Ca^2+^in mitochondria, since mitochondrial Ca^2+^ activates the electron transportation chain (Glancy et al., 2013). We attempted to buffer cytosolic Ca^2+^ using BAPTA to determine whether elevated cytosolic Ca^2+^ is responsible for the chemotaxis defects in MFN2 knockdown cells. However, a global cytosolic Ca^2+^ inhibition abrogated the ability of dHL-60 to migrate (Supplementary Fig. 6a, b), possibly due to the requirement of precisely regulated cytosolic Ca^2+^, spatially and/or temporally, for neutrophil migration (Mandeville and Maxfield, 1997; Marks and Maxfield, 1990). The mitochondrial calcium uniporter (MCU) is one of the primary sources of mitochondrial uptake of calcium which regulates migration of many cell types including primary human neutrophils (Zheng et al., 2017). The MCU inhibitor Ru360 did not cause further reduction of chemotaxis in MFN2 knockdown dHL-60 cells (Supplementary Fig. 6c, d and Supplementary Movie. 12), indicating that MCU and MFN2 lies in the same pathway in terms of regulating chemotaxis in dHL-60 cells.

**Figure 6.**
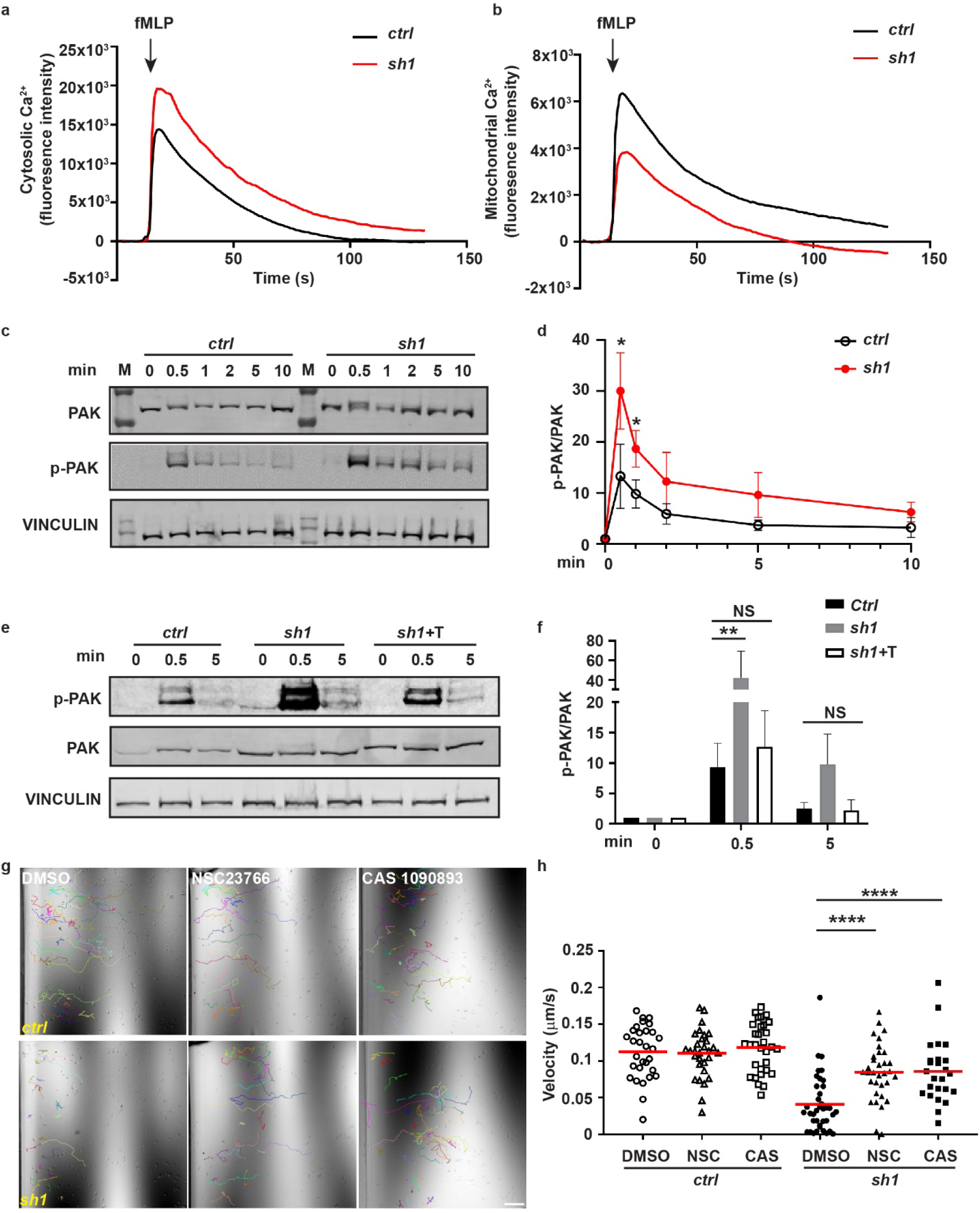
Mitochondria-ER tether restores RAC signaling and neutrophil chemotaxis in MFN2-deficient HL-60 cells. a) Cytosolic Ca^2+^ in the control or MFN2 knockdown cell lines after fMLP stimulation. b) Mitochondrial Ca^2+^ in the control or MFN2 knockdown cell lines after fMLP stimulation. c) Western blot determining the amount of pPAK in dHL-60 treated with fMLP at indicated time points. d) Quantification of p-PAK in different cell lines at indicated time points after fMLP stimulation a). e) Western blot determining the amount of pPAK in dHL-60 treated with fMLP at indicated time points and f) quantification of p-PAK in different cell lines at indicated time points after fMLP stimulation. g) Quantification of velocity of neutrophil chemotaxis to fMLP in the presence of vehicle or the Rac inhibitor NSC23766 or CAS1090893. h) Representative images with individual tracks. One representative result of three biological repeats is shown in a, b, c, e, g and h. Data are pooled from three independent experiments in d and f. n>20 cells were tracked or quantified in h. *, p<0.05, unpaired *t*-test in d; **, p<0.01; ****, p<0.0001, NS, no significance, one-way ANOVA (f and h). Scale bar: 100 µm in g.

Notably, the predominant cortical actin, nascent focal contacts and extensive membrane ruffles in *Mfn2-null* MEF cells (Fig. 3) resembled the classic phenotype of fibroblasts expressing the constitutively active Rac (Hall, 1998), indicating that Rac may be over-activated in Mfn2-depleted cells. Rac activation follows different time courses when cells are in suspension and when adhering to substrates (He et al., 2013). To measure Rac activation, we plated the cells on substrate-coated plates and used the phosphorylation of PAK (Graziano et al., 2017) as a readout. The phosphorylation of PAK peaked at 30 s post stimulation and returned to the baseline at 5 m post stimulation in control cells. Whereas in the MFN2 deficient cells, the phosphorylation of PAK were elevated at all the time points investigated compared with the control (Fig. 6c, d), suggesting a suppressive role of MFN2 in RAC activation in dHL-60 cells. Notably, the excessive phosphorylation of PAK can be corrected by expressing the artificial tether in MFN2-deficient dHL-60 cells (Fig. 6e, f), indicating that MFN2 moderates RAC activation via mediating mitochondria-ER tether. Furthermore, two different RAC inhibitors, NSC23766 or CAS1090893, restored neutrophil migration in *MFN2-sh1* cells, at least partially, at a concentration that did not affect chemotaxis of control cells (Fig. 6g, h and Supplementary Movie 13), further confirming that MFN2 regulates neutrophil migration through suppressing RAC activation. Together, Mfn2 mediated mitochondria-ER tether regulates the mitochondrial calcium uptake and RAC signaling to orchestrate chemotaxis in dHL-60 cells.

## Discussion

Here we report that Mfn2 is crucial for neutrophil adhesion and migration, providing evidence that Mfn2 regulates the actin cytoskeleton and cell migration. By maintaining the tether between the mitochondria and ER, Mfn2 orchestrates intracellular Ca^2+^ signaling and regulates Rac activation. Therefore, we have identified the mechanism for how Mfn2 regulates neutrophil adhesive migration, and highlighted the importance of mitochondria and their contact with the ER in neutrophils.

Mfn1 and Mfn2 possess unique functions, although both mediate mitochondrial outer membrane fusion (Chen et al., 2003). Our data that only MFN2, but not MFN1, is required for neutrophil adhesive migration supports that mitochondrial fusion is not likely important for neutrophil migration. Our observation goes against a body of literature that mitochondrial fission promotes cell migration in other cell types (Campello et al., 2006; Zhao et al., 2013). Alternatively, we propose a model that, in neutrophils, the interaction of mitochondria with the ER is more critical than a fused network. Although MFN2 is not the only protein that can mediate the mitochondria-ER tether (Eisenberg-Bord et al., 2016), mitochondria and ER interaction was significantly reduced upon MFN2 deletion in dHL-60, suggesting that MFN2 is at least one of the critical tether proteins in neutrophils. The fused mitochondrial network in neutrophils is possibly a result of the abundant expression of the mitofusins. Mutations in human MFN2 cause Charcot-Marie-Tooth disease type 2A (CMT2A), a classical axonal peripheral sensorimotor neuropathy (Zuchner et al., 2004). MFN2 is also implicated in many other diseases such as cancer, cardiomyopathies, diabetes and Alzheimer’s disease (Filadi et al., 2018). Currently, over 100 dominant mutations in the *MFN2* gene have been reported in CMT2A patients though how these mutations lead to disease is still largely unknown. The challenges in MFN2 research are that MFN2 regulates mitochondrial fusion and a plethora of cellular functions such as mitochondrial dynamics, transport, mtDNA stability, lipid metabolism and survival (Chandhok et al., 2018). In addition, gain-of-function and loss-of-function mutations are reported that affect different aspects of cellular functions (Chandhok et al., 2018). Our findings provide a new direction to understand the consequences of MFN2 deficiency in disease pathology, namely the actin cytoskeleton and Rac. Our findings also imply a possibility that the defects in immune cell migration in humans may affect immunity or chronic inflammation and indirectly regulate the progression of the aforementioned diseases. Future work will be required to carefully evaluate the individual mutations of MFN2 identified in human diseases in immune cell migration. It is possible that mutations disrupting mitochondria-ER tether, but not membrane fusion, result in defects in cell adhesion and the cytoskeleton regulation.

Our conclusions present a significant departure from the prevailing focus of bioenergy, in other word ATP, in cell migration. In many cell types, including neutrophils, the relevance of mitochondria-derived ATP in cell migration is emphasized (Bao et al., 2015; Bao et al., 2014). A recent report has confirmed the established literature that mitochondria do not provide ATP in neutrophils (Amini et al., 2018). Intriguingly, OPA1 deletion suppresses the production of neutrophil extracellular traps and alters the cellular ATP levels by indirectly suppressing glycolysis. In contrast, MFN2 deletion does not affect ATP levels (Supplementary Fig. 6) or affect neutrophil extracellular trap formation (Amini et al., 2018), suggesting again distinct biological functions of OPA1 and MFN2. In vascular endothelial cells, mitochondria also serve as signaling rather than energy-producing moieties (Lugus et al., 2011). In our study, in addition to the altered Ca^2+^ level, mitochondrial membrane potential and ROS, both critical for neutrophil chemotaxis and migration (Fossati et al., 2003; Zhou et al., 2018a), were reduced in stimulated *MFN2*-deficient dHL-60 cells. It remains to be determined whether the modest decreases in mitochondria membrane potential or ROS contribute to the defect of neutrophil migration upon MFN2 depletion.

Our result in leukocyte is consistent with previous results in murine fibroblasts (de Brito and Scorrano, 2008) that knocking out Mfn2 results in excessive cytosolic Ca^2+^ and defective mitochondrial calcium uptake. Intriguingly, chronic blockade of mitochondrial calcium import by depleting the mitochondrial calcium uniporter resulted in the reduction of the ER and cytosolic Ca^2+^ pools and a migration defect (Prudent et al., 2016). Although the phenotype is similar with MFN2 depletion, the underlying mechanisms is possibly different. Whereas MFN2 reduces the cytosolic Ca^2+^ levels after the activation of the chemokine receptor, MCU is required for maintaining the Ca^2+^ store and elevates the Ca^2+^, suggesting a requirement of delicate and precise calcium signaling in orchestrating neutrophil migration. Although cytosolic Ca^2+^ triggers the activation of Rac in slow moving cells (Price et al., 2003), previous work in neutrophils suggests that Rac activation was independent of cytosolic Ca^2+^ (Geijsen et al., 1999). This discrepancy could be explained with the differences in assay conditions (suspension vs. adhesion) or how Ca^2+^ levels are manipulated in experiments (elevation vs. reduction). Further work will be required to determine whether/how elevated calcium regulates Rac activation in neutrophils.

In summary, combining evidence from different models, we have discovered an essential role Mfn2 plays in neutrophil adhesion and migration, and determined the downstream mechanism, which provides insights and potential therapeutic strategies for inflammatory diseases and mitochondrial diseases.

## Methods

### Animals

The zebrafish and mice experiments were conducted in accordance to the internationally accepted standards. The Animal Care and Use Protocols were approved by The Purdue Animal Care and Use Committee (PACUC), adhering to the Guidelines for Use of Zebrafish and Mice in the NIH Intramural Research Program (Protocol number: 1401001018 and 1803001702). To generate transgenic zebrafish lines, plasmids with the tol2 backbone were coinjected with Tol2 transposase mRNA into embryos of the AB strain at one-cell stage as described (Zhou et al., 2018a). Constructs for neutrophil-specific knockout in zebrafish were generated as described (Zhou et al., 2018a) using the following primers (guide RNA sequences were indicated with underscores):

mfn2 guide1-F1: <u>GTGGATGAGCTGCGGGTGG</u>GTTTAAGAGCTATGCTGGAAACAGCATAGC; mfn2 guide1-R1: <u>CGCACCTCCGCCACCTGCC</u>CGAACTAGGAGCCTGGAGAACTGC; mfn2 guide1-F2: GGTGGCGGAGGTGCGGTTTAAGAGCTATGCTGGAAACAGCATAGC; mfn2 guide1-R2: CCGCAGCTCATCCACCGAACCAAGAGCTGGAGGGAGA; mfn2 guide2-F1: <u>GGGGGATACCTGTCCAAAG</u>GTTTAAGAGCTATGCTGGAAACAGCATAGCAAG; mfn2 guide2-R1: <u>AGACCTTCCTCTATGTGCC</u>CGAACTAGGAGCCTGGAGAACTGCTATATAAAC; mfn2 guide2-F2: CATAGAGGAAGGTCTGTTTAAGAGCTATGCTGGAAACAGCATAGCAAGTTTAAATAAG; mfn2 guide2-R2: F1: <u>GTAGTTGGGGACCAGAGTG</u>GTTTAAGAGCTATGCTGGAAACAGCATAGC; opa1 guide-R1: <u>CCTCACTGCTCAGCTGCC</u>CGAACTAGGAGCCTGGAGAACTGC; opa1 guide-F2: AGCTGAGCAGTGAGGGTTTAAGAGCTATGCTGGAAACAGCATAGC; opa1 guide-R2: CTGGTCCCCAACTACCGAACCAAGAGCTGGAGGGAGA.

All mice used in this study were purchased from Jackson Laboratories. Conditional *Mfn2* knockout mice (B6.129(Cg)-*Mfn2^tm3Dcc^*/J) were crossed to S100A8-Cre (B6.Cg-Tg(S100A8-cre, -EGFP)1Ilw/J) transgenic mice to obtain a homozygous floxed *Mfn2* alleles with or without the Cre. All mice were used at age 6-12 weeks, and both male and female were used for experiments.

### Cell Culture

The HL-60 line was a generous gift from Dr. Orion D. Weiner (UCSF). HEK293T, *wild-type*, *Mfn2-null* and *Mfn1-null* MEFs were purchased from American Type Culture Collection (ATCC). HUVEC was from Sigma-Aldrich (200P-05N). All cells were maintained at 37°C with 5% CO_2_. HL-60 cells were cultured in RPMI-1640 (Corning) with 10% FBS (Minipore), 25 mM HEPES, 1% sodium bicarbonate, and 1% sodium pyruvate. HEK293T and MEF cells were cultured in 10% FBS, 4.5g/glucose DMEM (Corning) with sodium bicarbonate. HUVEC was cultured in Endothelial Cell Growth Media (R&D SYSTEMS, CCM027). HL-60 cells were differentiated with 1.5% DMSO for 6 days. Cells were checked monthly for mycoplasma using the e-Myco plus Mycoplasma PCR Detection Kit (Bulldog Bio 25234).

To generate knockdown lines in HL-60 cells, pLKO.1 lentiviral constructs with shRNA was obtained from Sigma-Aldrich (*MFN2-sh1*:TRCN0000082684, *MFN2-sh2*: TRCN0000082687, *MFN1-sh*: TRCN0000051837), and SHC 003 was used as a non-targeting control. MFN2 rescue construct was generated by replacing GFP in TRCN0000082684 with *sh1*-resistant *MFN2*. Primers MFN2r-F: CAAGTGTATTGTGAAGAGATGCGTGAAGAGCGGCAAG and MFN2r-R: TTCACAATACACTTGTTGCTCCCGAGCCGCCATG was used to make *sh1*-resistant *MFN2* with Mfn2-YFP (addgene #28010) as the template. Primers: pLKO-F: AATTCTCGACCTCGAGACAAATGGC and pLKO-R: GGTGGCGACCGGGAGCGC were used to linearize the backbone of pLKO, and p-MFN2r-F: CTCCCGGTCGCCACCATGTCCCTGCTCTTCTCTCG and p-MFN2r-R: TCGAGGTCGAGAATTTTATCTGCTGGGCTGCAGGT were used to amplify *sh1*-resistant *MFN2* fragment. In-Fusion cloning (In-Fusion HD Cloning Plus kit, Clontech) was used to fuse the *sh1*-resistant *MFN2* fragment with the linearized backbone. Tether rescue construct was generated by replacing the GFP in TRCN0000082684 with a GFP containing Mitochondria localization signal (ATGGCAATCCAGTTGCGTTCGCTCTTCCCCTTGGCATTGCCCGGAATGCTGGCCCTCCTTGGCTGGTGGTGGTTTTTCTCTCGTAAAAAA) and ER localization signal (ATGGTTTATATTGGCATCGCTATTTTTTTGTTTTTGGTGGGCCTGTTTATGAAA) at it N- and C-terminal respectively. Primers: Tether rescue+: CTCCCGGTCGCCACC ATGGCAATCCAGTTGCGTTCG, Tether rescue-: TCGAGGTCGAGAATTTTAAGATACATTGATGAGTTTGG. Transomics shERWOOD UltramiR lentiviral inducible shRNA system (non-targeting control: TLNSU4300, mfn2 shRNA: ULTRA-3418270) were used for acute MFN2 deletion. Lentiviral constructs together with pCMV-dR8.2 dvpr (addgene #8455) and pCMV-VSV-G (addgene #8454) were co-transfected into HEK293T cells with Lipofectamin 3000 (Invitrogen L3000015) to produce lentivirus. Virus supernatant was collected at both 48 hpt and 72 hpt, and further concentrated with Lenti-X concentrator (Clotech 631232). HL-60 cells were infected in complete medium supplemented with 4 µg/ml polybrene (Sigma TR-1003-G) and selected with 1 µg/ml puromycin (Gibco A1113803) to generate stable line.

### Microinjection

Microinjections of fish embryos were performed as described (Deng et al., 2011). Briefly, 1 nl of mixture containing 25 ng/µl plasmid and 35 ng/µl Tol2 transposase mRNA was injected into the cytoplasm of embryos at one-cell stage.

### Tailfin wounding and Sudan black staining

Tailfin wounding and Sudan Black staining were carried out with 3 dpf embryos as described (Zhou et al., 2018b). Briefly, embryos were fixed in 4% paraformaldehyde in phosphate-buffered saline overnight at 4°C and stained with Sudan black.

### Live imaging

Time-lapse images for zebrafish circulation, LTB_4_ bath, flow adhesion assay were obtained with AXIO Zoom V16 microscope (Zeiss). Time-lapse fluorescence images for zebrafish neutrophil motility were acquired using a laser scanning confocal microscope (LSM 710, Zeiss) with a 1.0/20 x objective lens at 1 min interval of 30 min. Neutrophils were tracked using ImageJ with MTrackJ plugin and the velocity was plotted in Prism 6.0 (GraphPad). Time-lapse fluorescence images for dHL-60 migration were acquired using a laser scanning confocal microscope (LSM 710, Zeiss) with a 1.0/20 x objective lens at 10 sec interval for 5 min. Cells were stained with 1 µM ER-tracker (Invitrogen E34251) and 20 nM TMRM (Invitrogen T668) for 20 min, washed twice with HBSS and added to fibrinogen coated wells. After 30 min, cells were treated with 1 nM fMLP to induce chemokinesis.

### Confocal imaging

For confocal imaging, images were obtained using a laser-scanning confocal microscope (LSM 800, Zeiss) with a 1.4/63x oil immersion objective lens. Images were analysis with ImageJ. For fluorescence intensity measurement, images within an experiment were acquired using identical camera settings and background was subtracted using ImageJ with the rolling ball radius of 50. Mean fluorescence intensity of selected areas was measured by Measurement in ImageJ and plotted in Prism (GraphPad). Colocalization was using ImageJ Plugin Coloc 2. Interaction between channels was quantified by Manders’ colocalization coefficient as described (de Brito and Scorrano, 2008).

### µ-slide Chemotaxis

Differentiated HL-60 cells were resuspended in mHBSS at 4×10^6^/ml and loaded into µ-slides (ibidi, 80322) following manufacturer’s instructions. fMLP was added to the right reservoir at a concentration of 1 µM. Chemotaxis was recorded every 1 min for 2 h using a laser scanning confocal microscope (LSM 710, Zeiss) with a 1.0/10 x objective. The velocity of neutrophils was measured using ImageJ with MTrackJ plugin and plotted in Prism 6.0 (GraphPad). For inhibitor treatments, dHL-cells were pre-treated with DMSO, NSC23766 (200 µM, Sigma SML0952), AIP (50 µM, R&D SYSTEMS 5959/1), CAS 1090893 (50 µM, Millipore 553511), or RU360 (50 µM, Millipore 557440) for 30 min before loading into µ-slides. For 3D migration, dHL-60 cells were starved for 1 h in HBSS with 0.1% FBS and 20 mM HEPES. 100 µl of cells at 3×10^6^ cells/ml were added to 33 µl of Matrigel (Corning #356235), mixed gently and loaded in µ-slides (ibidi, 80322). The slide was incubated for 15 min at 37 °C to allow matrix solidification. fMLP at 187.5nM was added to the right reservoir. Chemotaxis was recorded every 1 min for 2 h, with a Lionheart FX Automated Microscope (Biotek) using a 10x phase objective, Plan Fluorite WD 10 NA 0.3 (1320516). Cells were tracked using MTrackJ image J plugin and plotted in Prism 6.0 (GraphPad).

### Flow adhesion

Neutrophil flow adhesion assay was performed as described (Zhou et al., 2014). Briefly, 5×10^5^ HUVEC cells in 2 ml were plated onto 10 µg/ml fibrinogen-coated 35 mm plate (Corning 430165), and incubated at 37°C. Then the HUVEC monolayer was primed by 20 ng/ml human TNF-a (Life technologies PHC3015) for 4-6 h. dHL-60 cells were harvested and resuspended at a cell density of 5×10^5^ cells/ml in complete medium. dHL-60 cells were flowed on top of HUVEC monolayer at a speed of 350 µl/min using a syringe pump. Cells adhering to the monolayer were recorded using AXIO Zoom V16 microscope (Zeiss) with camera streaming for 5 min. The total number of adherent neutrophils were quantified at 5 min.

### Western Blot

Protein samples were separated using SDS-PAGE and transferred onto nitrocellulose membranes (LI-COR NC9680617). Membranes were blocked for ∼30 min in PBST (PBS and 0.1% Tween 20) with 5% BSA. After blocking, membranes were incubated O/N with primary antibodies diluted 1:1,000 in PBST at 4°C and secondary antibodies diluted 1:10,000 in PBST at room temperature for 1 h. Odyssey (LI-Cor) was used to image membranes. Primary antibodies anti-Mfn2 (Cell Signaling 9482S), anti-Mfn1 (Cell Signaling 14793S), anti-beta-Tubulin (DSHB, E7), anti-Rac1/2/3 (Cell Signaling 2465S) and secondary antibodies anti-rabbit (ThermoFisher SA5-35571), anti-mouse (Invitrogen 35518) were used. Phospho-PAK level was determined as described (Graziano et al., 2017). 1∼2×10^6^ dHL-60 cells were adhered to fibrinogen coated 6cm dishes for 30 min, and stimulated with 100nM fMLP for indicated time. Ice cold stop solution (20% TCA, 40 mM NaF, and 20 mM β-glycerophosphate, (Sigma #T6399, #G9422, #s6776)) was immediately added to the cells at 1:1 volume and put on ice for 1h. Lysates were pelleted and washed once with 0.75 ml of ice cold 0.5% TCA and resuspended in 2× Laemmli sample buffer (Bio-Rad Laboratories). Western blot was performed as described (Hsu et al., 2019) with anti-phospho-PAK (Cell signaling #2605S), anti-PAK (Cell signaling #2604) along with Vinculin (Sigma #V9193) as a loading control.

### Bone Marrow Neutrophil isolation

Femurs and Tibias from mice 8-12 weeks of age were isolated and whole bone marrow was isolated, and passed through a 70 µm filter followed by RBC lysis (Qiagen 158904). Bone marrow neutrophils were isolated using a negative selection column (MACS 130-097-658). Neutrophils viability was determined by trypan blue staining showing >99% viability.

### Peritonitis model

1ml of 4% thioglycollate (Sigma B2551) was injected directly into the peritoneal cavity of mice 6-8 weeks of age. After 3 hours of incubation, peritoneal ascites were collected by introducing 8 ml of PBS into the cavity and collecting the ascites immediately afterwards. Cells were subjected to RBC lysis and viability was determined using trypan blue staining. Cells were stained with CD11b (BD 557686) and Ly6G (BD 566453) on ice for 30 minutes and washed 3 times with staining buffer. Cells profiles were collected with a BD fortessa analyzer and analyzed with Beckman kaluza software. Neutrophil population was defined as FCS/SSC high and CD11b+Ly6G^high^. Percentage of neutrophils in the lavage relative to total viable cells in each experiment was normalized to the sex-matched littermate control.

### Immunostaining

dHL-60 cells were resuspended in mHBSS and attached to fibrinogen-coated slides for 30 min. Then cells were stimulated with 100 nM fMLP for 3 min and fixed with 3% paraformaldehyde in PBS for 15 min at 37°C. The immunostaining of fixed cells were performed as described (Fayngerts et al., 2017). Briefly, after fixation, cells were permeabilized in PBS with 0.1% Triton X-100 and 3% BSA for 1 h at room temperature. dHL-60 cells were incubated with phalloidin - AlexaFluor 488 (Invitrogen A12379) or primary antibodies diluted 1:100 in 3% BSA overnight at 4 °C. The cells were then stained with secondary antibodies diluted 1:500 in 3% BSA and DAPI (Invitrogen D3571) for 1 h at room temperature. For MEF staining, cells were plated on fibrinogen-coated slides and incubated for ∼4 h at 37 °C, followed with fixation with 3% paraformaldehyde in PBS. Primary antibodies anti-Mfn2 (Cell Signaling 9482S), anti-TOMM20 (Santa Cruz sc-17764), anti-Calnexin (Cell Signaling 2433S), anti-Tubulin (Sigma T5168), anti-Paxillin (Invitrogen AHO0492), and secondary antibodis anti-rabbbit AlexaFluor 568 (Invitrogen A-11011), anti-mouse AlexaFluor 647 (Invitrogen A21236) were used.

### Electron microscopy

Transmission Electron Microscopy was performed at Purdue Life Science Microscopy Facility. dHL-60 cells were pelleted and fixed in 2.5% glutaraldehyde in 0.1 M sodium cacodylate buffer, post-fixed in buffered 1% osmium tetroxide containing 0.8% potassium ferricyanide, and en bloc stained in 1% aqueous uranyl acetate. They were then dehydrated with a graded series of acetonitrile and embedded in EMbed-812 resin. Thin sections (80nm) were cut on a Leica EM UC6 ultramicrotome and stained with 2% uranyl acetate and lead citrate. Images were acquired using a Gatan US1000 2K CCD camera on a FEI Tecnai G2 20 electron microscope equipped with a LaB6 source and operating at 100 kV or 200kV.

### Ca^2+^ measurement

Fluo-4 Calcium Imaging Kit (Invitrogen F10489) was used for cytosolic Ca^2+^ measurement. dHL-60 cells were resuspended in mHBSS and incubated with PowerLoad solution and Fluo-4 dye at 37 °C for 15 min and then at room temperature for 15 min. After incubation, cells were washed with mHBSS and loaded into fibrinogen-coated 96-well plates (greiner bio-one 655069) with 20,000 cells in 150 ul for each well, followed by incubation at 37 °C for 30 min. Green fluorescence images were recorded by BioTek Lionheart FX Automated Microscope with 20x phase lens at 1s interval of 10s. Then 15 ul of 1 uM fMLP was injected into cells using the injector of BioTek Lionheart FX Automated Microscope. Images were recorded for another 2 min with 1s interval. The fluorescence intensity of basal Ca^2+^ level was set to 0.

For mitochondrial Ca^2+^ measurement, Rhod-2 (Invitrogen R1245MP) was used. dHL-60 cells were incubated in mHBSS with Rhod-2 at 37 °C for 30 min, and then washed and added into fibrinogen-coated 96-well plates with 150 µl/well. After 30 min incubation, time-lapse red fluorescence images were acquired by BioTek Lionheart FX Automated Microscope with 1s interval of 10s and followed by fMLP injection and image recording for another 2 min with 1s interval. The fluorescence intensity of basal Ca^2+^ was normalized to 0.

### Cell spreading

MEF cell spreading assay was performed as described (Jovic et al., 2007). Briefly, cells were trypsinized and replated onto fibrinogen-coated µ-slide 8 well plates (ibidi 80826) with complete medium. Time-lapse images were acquired using BioTek Lionheart FX Automated Microscope with 20x phase lens at 2 min interval of ∼3 h at 37°C with 5% CO_2_.

### Wound closure

MEF cells in complete medium were seeded onto 96-well plates (FALCON 353075) and incubated at 37 °C overnight. A wound was induced by automated 96-well WoundScratcher (BioTek) for each well. Cells were washed twice with mHBSS and time-lapse images were acquired using BioTek Lionheart FX Automated Microscope with 4x phase lens at 20 min interval of ∼12 h at 37°C with 5% CO_2_.

### Flow cytometry analysis

dHL-60 cells were harvested and resuspended into ice-cold FACS buffer (PBS with 1% BSA) at a concentration of 1×10^6^ cells/ml. 5 ul of Annexin V (BD 563973) solution were added into 100 ul cell suspension and incubated at room temperature for 30 min. Cells were washed for three times with ice-cold FACS buffer, and followed by flow cytometry analysis. For surface markers, cells were incubated on ice for 1 hour in staining buffer (1% BSA in PBS) containing CD18-PE (BD #B555924), PE isotype control (BD #B554680), CD11b-AF647 (Biolegend #301319), AF647 isotype control (Biolegend # 400130), CD15-BV510 (BD #B563141), BV510 isotype control (BD #B562946) or WGA-AF594 (thermos #W11262), washed three times with staining buffer and resuspended in suitable volumes. Flow cytometry was performed using BD Fortessa analyzer.

### Quantitative RT-PCR

Neutrophils from *Mfn2* conditional KO mice were isolated and RNA was extracted using Qiagen RNeasy Mini Kit (74104). HL-60 cells were differentiated for 6 days in 1.3% DMSO and RNA was extracted using Qiagen RNeasy Mini Kit (74104). Messenger RNAs were reverse-transcribed with Transcriptor First Strand cDNA Synthesis Kit (Roche). qPCR were performed using the FastStart Essential DNA Green Master (Roche) in a LightCycler® 96 Real-Time PCR System (Roche Life Science). Primers: mus-mfn2+: 5’-tctttctgactccagccatgt-3’; mus-mfn2-: 5’-tggaacagaggagaagtttctagc-3’; mus-gapdh+: 5’-gggttcctataaatacggactgc-3’; mus-gapdh-:5’-ccattttgtctacgggacga-3’; hsa-mmp9+: 5’-gaaccaatctcaccgacagg-3’; hsa-mmp9-: 5’-gccacccgagtgtaaccata-3’; hsa-rpl32+:5’-gaagttcctggtccacaacg-3’; hsa-rpl32-:5’-gagcgatctcggcacagta-3’. The specificity of the primers was verified as a single peak in the melt-curves. The relative levels of mRNA were calculated using the deltaCt method. The relative fold change with correction of the primer efficiencies was calculated following instructions provided by Real-time PCR Miner (http://ewindup.info/miner/data_submit.htm) (Zhao and Fernald, 2005).

### Mitochondrial membrane potential, ROS, and ATP measurement

Mitochondrial membrane potential was measured using TMRM. dHL-60 cells were resuspended in mHBSS and incubated with 150 nM MitoTracker (Invitrogen M22426), 20 nM TMRM (Invitrogen T668), and 0.2 µg/ml Hoechst (Invitrogen H3570) for 30 min at 37 °C. Then cells were washed and plated onto fibrinogen-coated µ-slide 8 well plates. After 30 min incubation, cells were stimulated with or without fMLP at a concentration of 100 nM. The fluorescent images were acquired using BioTek Lionheart FX Automated Microscope with 20x phase lens, and processed using ImageJ. Mitochondrial membrane potential was measured using the fluorescence intensity of TMRM normalized to the intensity of MitoTracker of each cell. For mitochondrial ROS measurement, MitoSOX (M36008) was used. 5 µM of mitoROX was added to the cell suspension and incubated for 30 min at 37 °C. Cellular ATP level was measured using ATP Assay Kit (abcam #ab83355) according to the assay procedure. Briefly, dHL-60 cells treated with or without fMLP were harvested, washed with PBS, and resuspended in ATP Assay Buffer. Samples with ATP reaction mix were loaded into 96-well plate (greiner bio-one 655069) and incubated at room temperature for 30 min protected from light. Results were measured using a microplate reader (BioTek) at Ex/Em = 535/587. All results were normalized to control cell lines without fMLP treatment.

### Mutational Efficiency Quantification

The mutation efficiency of neutrophil-specific knockout in zebrafish was quantified as described (Zhou et al., 2018a). To determine the mutation efficiency in *Tg(lyzC:Cas9-mfn2 sgRNAs), Tg(lyzC:Cas9-mfn2 sgRNAs#2)*, and *Tg(lyzC:Cas9-opa1 sgRNAs),* 3 dpf embryos of each line were digested with trypsin to prepare single cell suspensions. mCherry positive cells were sorted by FACS. Genomic DNA was purified using QIAamp DNA Mini Kit (Qiagen) from sorted cells. 5 µg of poly(dA:dT) (Invivogen) were used as the carrier DNA. The *mfn2 and opa1* loci around the sgRNA binding sites were amplified using PCR with the following primers: mfn2-F1: GGCGATGATAAACATGGCAGTTTG, mfn2-R1: GTACCACAGGTGCACAGTGTC, mfn2-F2: CTGGGACGCATCGGCCAATG, mfn2-R2: CTACCTGCTTCAGGCATTCCCTG, mfn2#2-F1: GTCGGGCTTCTCCTAAGTTATTC, mfn2#2-R1: CAGTGTCCATAGCCTAGAGTCTGC, mfn2#2-F2: GTGGTCTCATATAATTTTGCTTGCTG, mfn2#2-R2: CACACGCGAATCGATAAGAGGAAT, opa1-F1: CAAGCTCATTAAAGGTTTGAAACCACTTG, opa1-R1: CTCCACAAATCACATAGGTGAC, opa1-F2: GTGCCTGAATGCTCTACACTTTC, opa1-R: CATGATAACAATACCATGCACATGC, Purified PCR products were used for library construction with Nextera library prep kit and sequenced using an Illumina MiSeq 300 at the sequencing center of Purdue University. Raw reads have been deposited to the Sequence Read Archive (Accession number PRJNA510381, https://www.ncbi.nlm.nih.gov/Traces/study/?acc=SRP132222). Mutational efficiency was calculated using the CrispRVariants R package (Lindsay et al., 2016).

### Statistical analysis

Statistical analysis was performed with Prism 6 (GraphPad). Two-tailed student’s *t* test, or ANOVA was used to determine the statistical significance of differences between groups. A *P* value of less than 0.05 indicated in the figures by asterisks was considered as statistically significant.

## Acknowledgements

The clonal HL-60 cell line was generated by Alba Diz-Munoz (EMBL Heidelberg) and a generous gift from the laboratory of Orion Weiner (UCSF). Mice work was initiated with the generous help from Dr. Yao Liu in the laboratory of Zhao-Qing Luo (Purdue). We thank our undergraduate students, Abby Pei Lemke, Peiyi Yu and Simon Tian for cell tracking, Weicheng Wang for assistance with co-localization analysis. The work was supported by National Institutes of Health [R35GM119787 to DQ], [R01HD073156 to DM] and [P30CA023168 to Purdue Center for Cancer Research] for shared resources. WZ is supported by Cagiantas Fellowship, Purdue University. AH is supported by Purdue Research Foundation Grant and Cagiantas Fellowship, Purdue University.

## Author contributions

WZ, MNS, DU and DQ designed research and wrote the manuscript. WZ and AH performed most experiments and analyzed the data. YW, TW, HM, XW helped with experiments. JJ helped with data analysis. All authors read and approved the manuscript.

## Data availability statement

Plasmids are available on Addgene: p3E-U6a-U6c-mfn2 guide (Plasmid #121993); p3E-U6a-U6c-mfn2 guide#2 (Plasmid #121994); p3E-U6a-U6c-opa1 guide (Plasmid #121995); MYs-IRES-mitoRFP (Plasmid #121996); pMYs-MFN2-IRES-mitoRFP (Plasmid #121997).

