## Supplemental Figures for "Mitofusin 2 regulates neutrophil adhesive migration and the actin cytoskeleton"

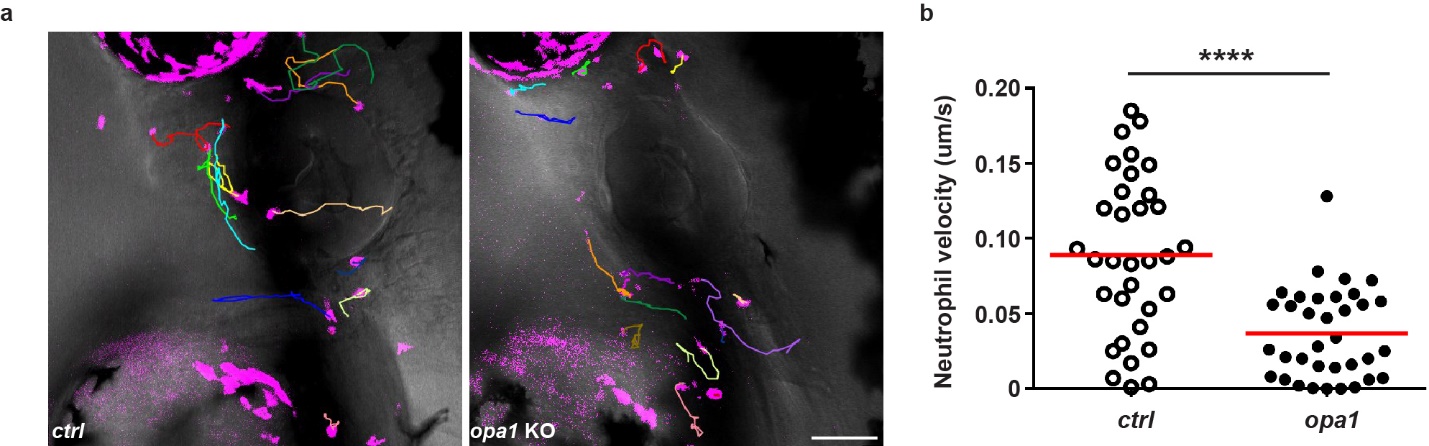
**Figure S1. Disrupting *opa1* reduces neutrophil motility in zebrafish.**

a) Representative tracks and b) Quantification of neutrophil motility in the head mesenchyme of the control line and the line with neutrophil specific *opa1* knockout at 3 dpf. One representative result of three biological repeats is shown. n>20 cells were tracked. ****, p<0.0001, unpaired *t* test. Scale bar: 50 µm.

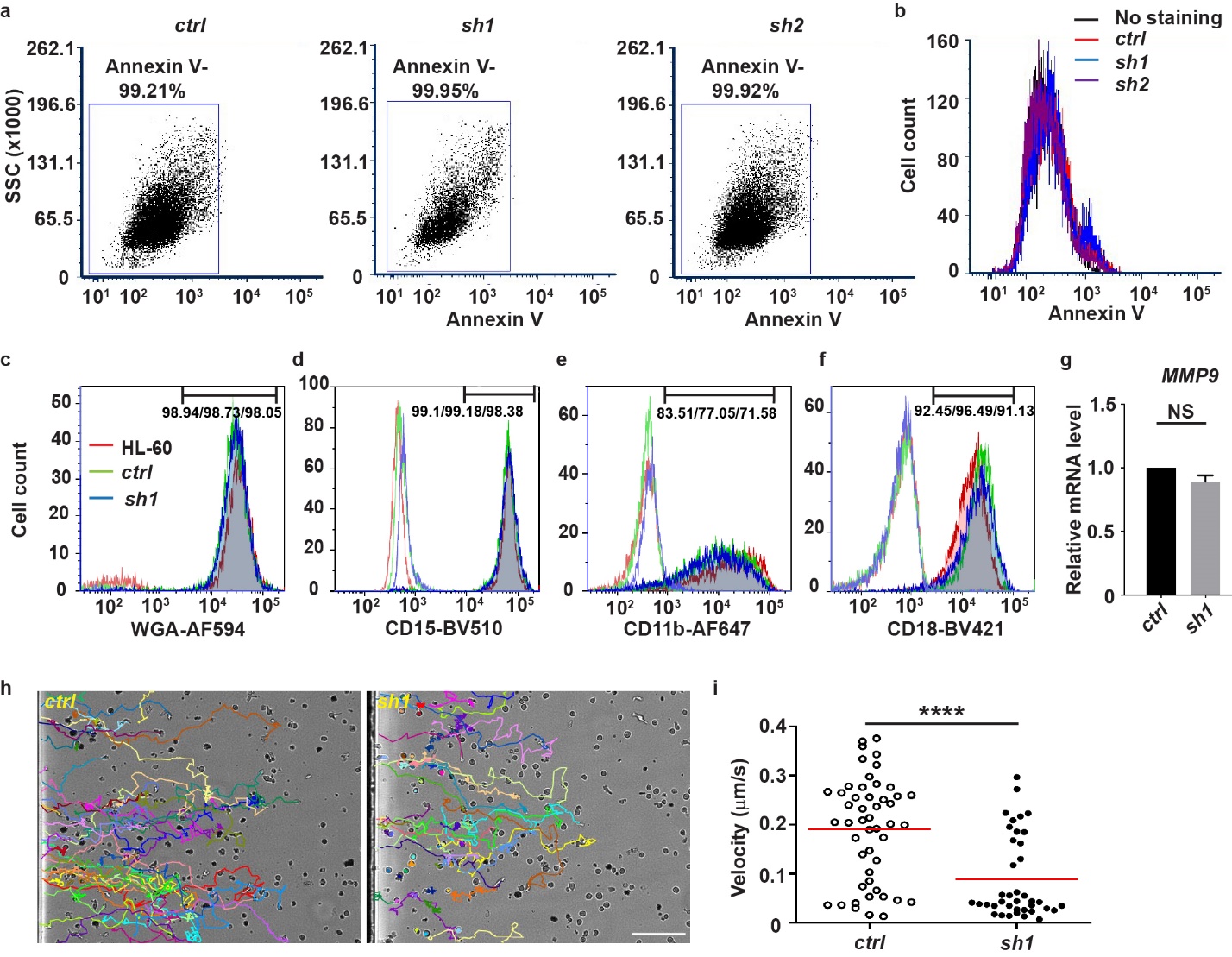
**Figure S2. MFN2 deficiency does not induce apoptosis or affect cell differentiation in dHL-60 cells.**

Apoptosis was determined by Annexin V staining on the surface and followed by flow cytometry in indicated cell lines. a) Dot plot and b) Histogram of Annexin V staining. c-f) Histogram of WGA, CD15, CD11b, and CD18 staining in indicated cell lines. g) Relative mRNA level of *MMP9* in indicated cells lines. One representative result of three biological repeats is shown (a-f). Data are pooled from three independent experiments in g.

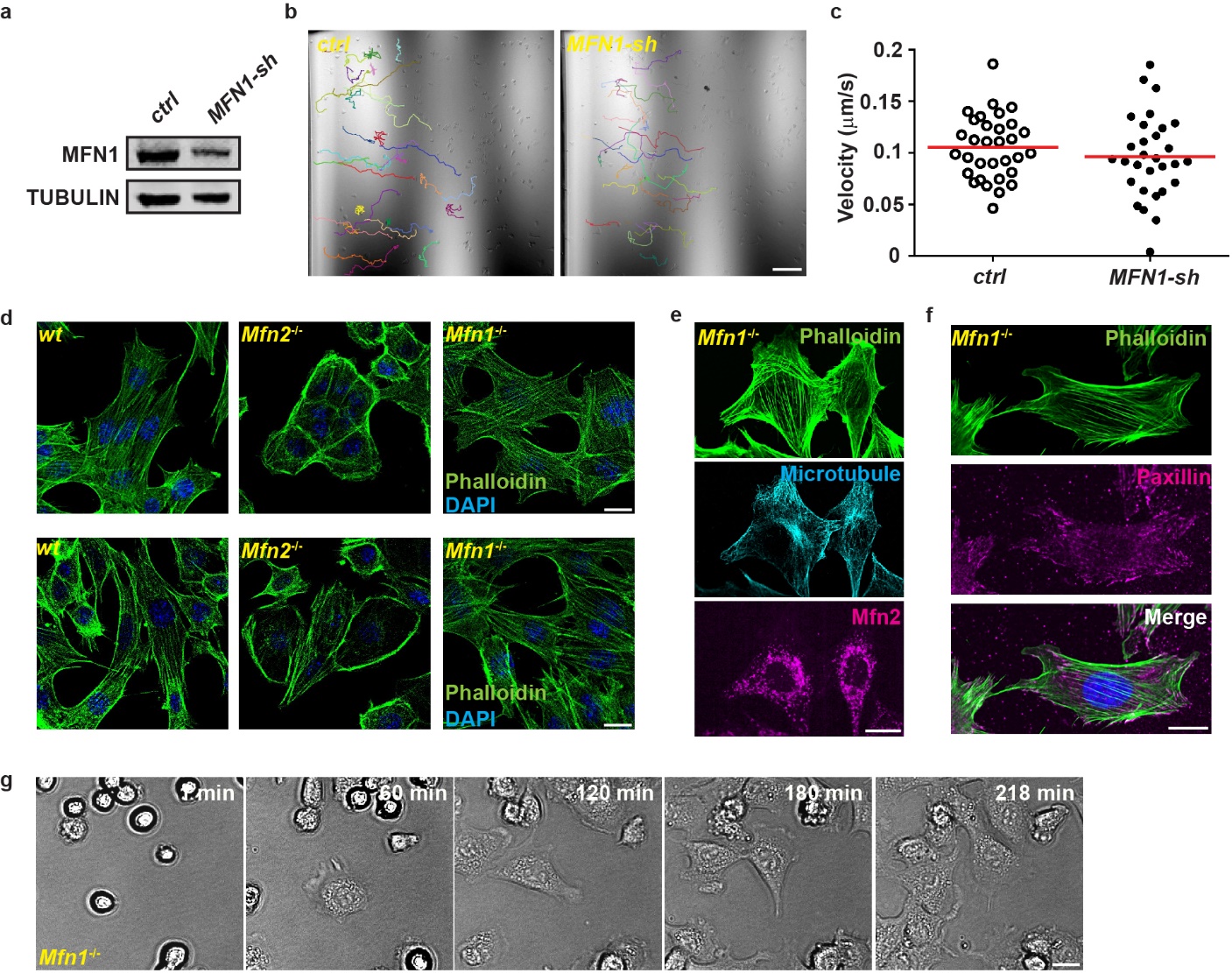

**Figure S3. MFN1 does not regulate adhesive migration of dHL-60 cells or actin cytoskeleton in MEF.** a)Western blot showing the expression level of MFN1 in the control cell line or the one stably expressing a shRNA targeting *MFN1*. b) Quantification and c) Representative tracks of dHL-60 cells migrating toward fMLP. d) Immunofluorescence of F-actin (Phalloidin) and nucleus (DAPI) in indicated cells plated on un-coated (upper) or fibrinogen-coated (lower) cover glasses. e) Immunofluorescence of F-actin (Phalloidin), microtubule and Mfn2 in *Mfn1-null* MEFs. f) Immunofluorescence of F-actin (Phalloidin) and Paxillin in *Mfn1-null* MEFs. g) Images of *Mfn1-null* MEFs spreading at indicated time points after plating. One representative result of three repeats is shown. n>20 cells were quantified and tracked in b and c. Scale bar: 100 µm in b, 10 µm in d-g.

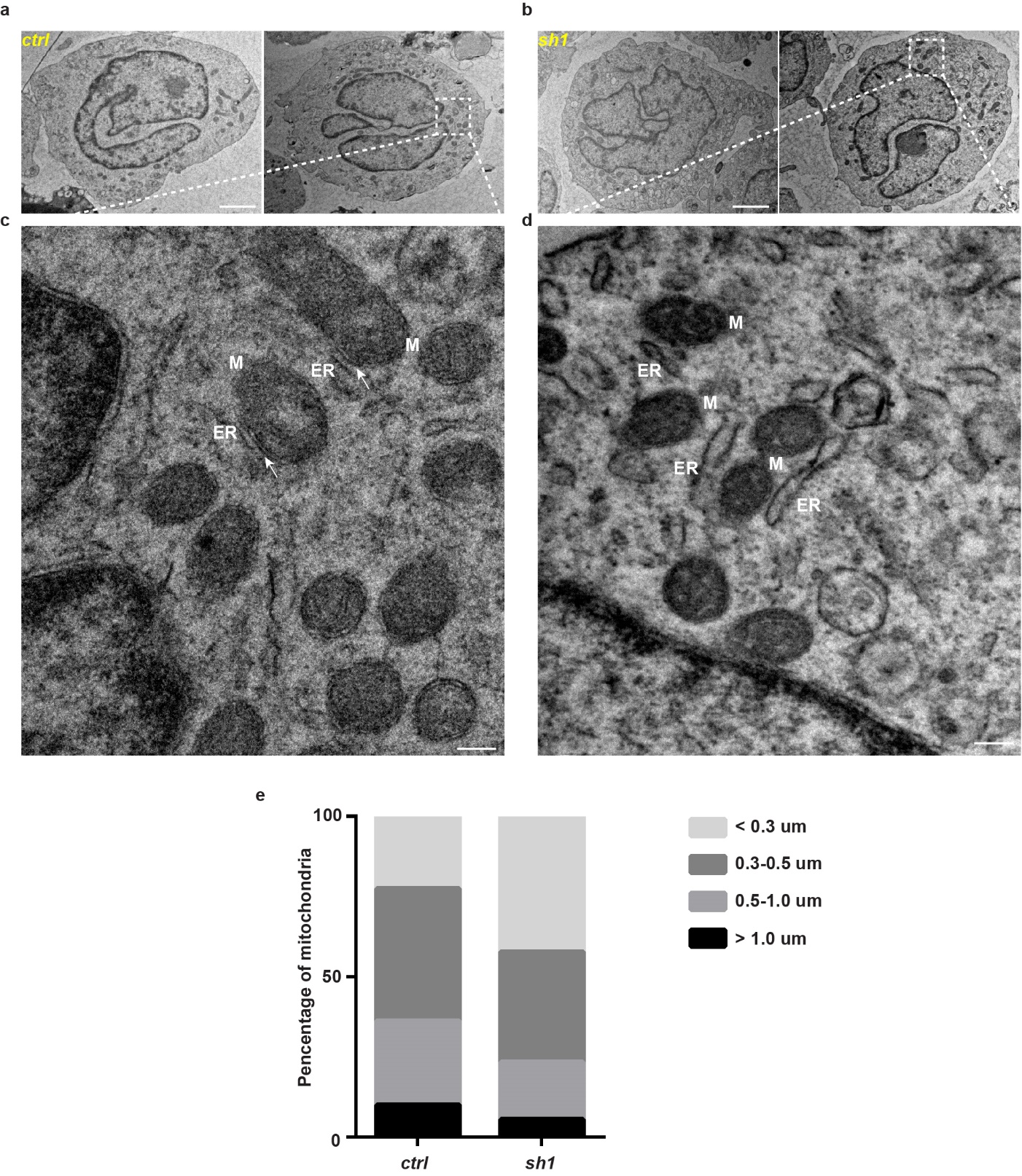

**Figure S4. Electron microscopy of mitochondria and ER in MFN2 knock down dHL-60 cells.** a, b) Representative electron microscopy images of control and MFN2 knockdown dHL-60 cells. c, d) Enlarged images of the white box in a) and b). M: mitochondria, ER: endoplasmic reticulum. White arrows: contact sites between mitochondria and ER. e) Quantification of the length of mitochondria in control and *MFN2-sh1* dHL-60 cells. n>20 cells were quantified in e. Scale bar: 2 µm in a and b; 0.2 µm in c and d.

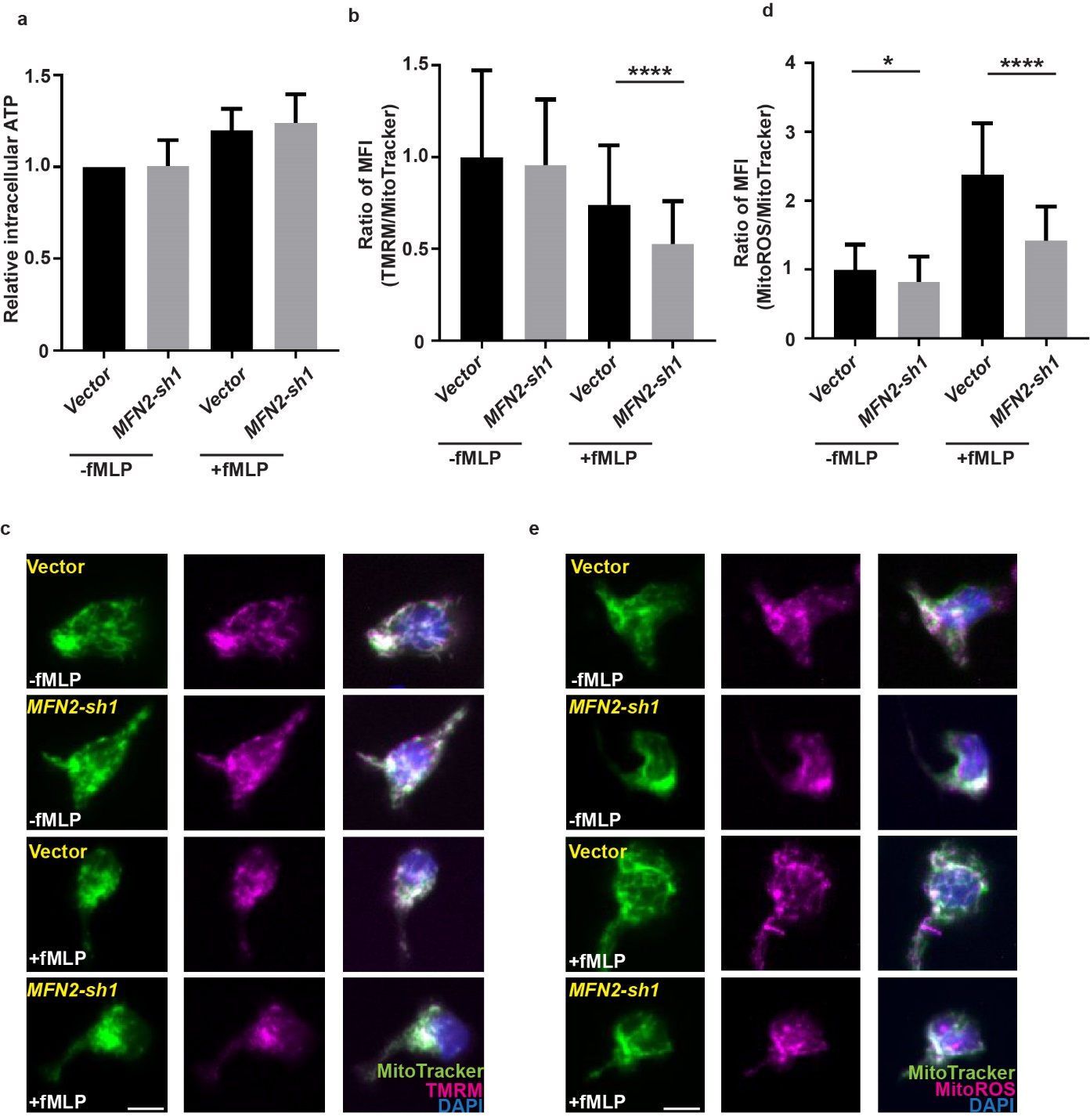

**Figure S5. MFN2 regulates mitochondrial membrane potential and ROS, but not ATP in dHL-60 cells.** a) Quantification of cellular ATP levels with or without treatment of fMLP for 3 min. b) Quantification of the mean fluorescence intensity of TMRM normalized with MitoTracker. c) Fluorescence of mitochondria (MitoTracker), membrane potential (TMRM), and nucleus (Hoechst) in the control or *MFN2* knockdown cell lines with or without treatment of fMLP for 3 min. d) Quantification of mean fluorescence intensity of MitoROS normalized with MitoTracker. e) Fluorescence of mitochondria (MitoTracker), mitochondrial ROS (MitoROS), and nucleus (Hoechst) in indicated cells with or without treatment of fMLP for 3 min. Data are pooled from three independent experiments (a, b, d) *, p<0.05; ****, p<0.0001, NS, no significance, by unpaired *t*-test. Scale bar: 10 µm in c and e.

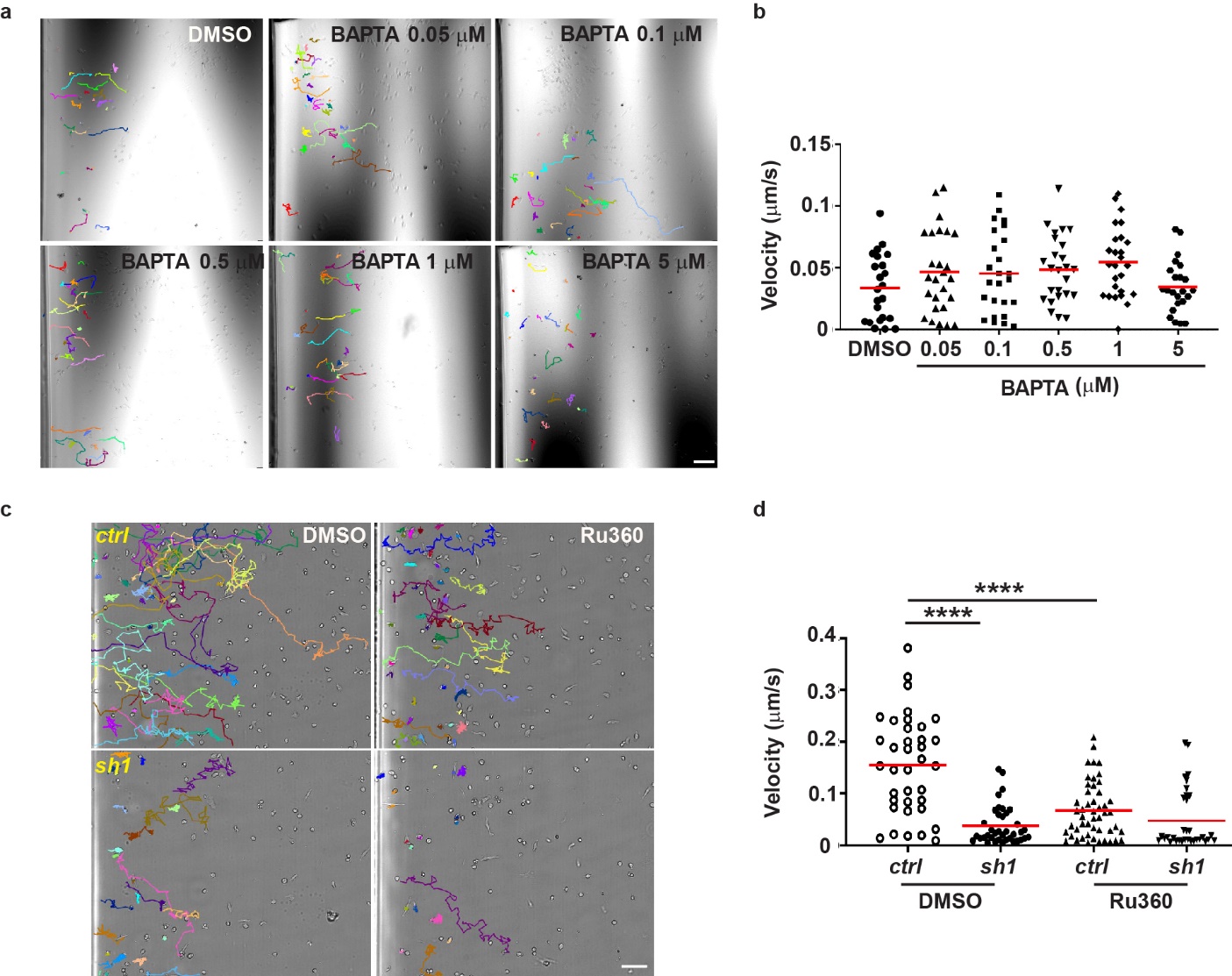

**Figure S6. Buffering intracellular Ca^2+^ does not rescue the migration defect of *MFN2* knockdown dHL-60 cells.**

a) Representative tracks and b) Quantification of MFN2 knockdown dHL-60 cells migrating toward fMLP in the presence of indicated concentrations of BAPTA. One representative result of three repeats is shown. c) Representative tracks and d) Quantification of neutrophil velocity migrating to fMLP in the presence of DMSO or MCU inhibitor, Ru360. One representative result of three repeats is shown in a-d. n>20 cells were quantified and tracked in a-d. ****, p<0.0001, by unpaired *t*-test in d. Scale bar: 100 µm.

**Supplementary Movies**

**Movie S1. Neutrophil specific deletion of *mfn2* alters neutrophil homeostasis in zebrafish.**

Time-lapse images of neutrophil homeostasis in 3 dpf embryos from the transgenic lines with neutrophil specific *mfn2* knockout and the control. Scale bar: 50 µm.

**Movie S2. Neutrophil specific deletion of *mfn2* using a second set of sgRNAs results in a similar phenotype.**

Time-lapse images of neutrophil homeostasis in 3 dpf embryos from a second transgenic line expressing different sgRNAs for neutrophil specific *mfn2* knockout. Scale bar: 50 µm.

**Movie S3. Opa1 is required for neutrophil motility in zebrafish.**

Time-lapse images of neutrophil migration at the head mesenchyme and individual neutrophils were tracked for velocity quantification. Scale bar: 50 µm.

**Movie S4. Mfn2 is required for neutrophil chemotaxis to LTB_4_ in zebrafish.**

Time-lapse images of neutrophil chemotaxis in 3 dpf embryos from the transgenic lines with neutrophil specific *mfn2* knockout and the control after LTB4 treatment. Scale bar: 50 µm.

**Movie S5. MFN2 regulates adhesive migration in dHL-60 cells.**

Chemotaxis of the control, *MFN2* knockdown, and *MFN2* rescue cells towards fMLP in the µ-slide. Individual neutrophils in the left half of the chamber were tracked. Scale bar: 50 µm.

**Movie S6. Acute deletion of MFN2 reduced adhesive migration in dHL-60 cells.**

Chemotaxis of the control with dox, *MFN2*-sh1 with or without dox cells towards fMLP in the µ-slide. Individual neutrophils in the left half of the chamber were tracked. Dox: doxycycline. Scale bar: 50 µm.

**Movie S7. *MFN2*-deficient dHL-60 cells have defective adhesion to endothelial cells.**

HUVEC cell monolayer was stimulated by 20 ng/ml TNF-a for 4-6 h. Vector control and *MFN2-sh1* dHL-60 cells were flowed through HUVEC layers with a speed of 350 ul/min and recorded by microscope with no interval. Scale bar: 50 µm.

**Movie S8. MFN1 does not regulate adhesive migration in dHL-60 cells.**

Chemotaxis of the control and *MFN1* knockdown cells towards fMLP in the µ-slide. Individual neutrophils in the left half of the chamber were tracked. Scale bar: 50 µm.

**Movie S9. MFN2 regulates wound closure of MEF cells.**

Wound closure of the control and *Mfn2^-/-^* MEFs. Scale bar: 200 µm.

**Movie S10. MFN2 regulates spreading of MEF cells.**

Spreading of the control, *Mfn2^-/-^* and *Mfn1^-/-^* cells on µ-slide 8 well plates immediately after plating. Scale bar: 20 µm.

**Movie S11. Restoring ER-mitochondrial tether rescue the migration defects in *MFN2*-deficient dHL-60 cells.**

Chemotaxis of control, *MFN2*-deficient, and *MFN2*-deficient cells with synthetic tether construct towards fMLP in the µ-slide. Individual neutrophils in the left half of the chamber were tracked. Scale bar: 50 µm.

**Movie S12. Inhibition of MCU does not further decrease chemotaxis in *MFN2*-deficient dHL-60 cells.**

Chemotaxis of control and *MFN2* knockdown cells treated with DMSO or Ru360 (50 µM) towards fMLP in the µ-slide. Individual neutrophils in the left half of the chamber were tracked. Scale bar: 50 µm.

**Movie S13. Rac inhibition restores chemotaxis in *MFN2*-deficient dHL-60 cells.**

Chemotaxis of control and *MFN2* knockdown cells treated with DMSO, NSC23766 (200 µM) or CAS 1090893 (50 µM) towards fMLP in the µ-slide. Individual neutrophils in the left half of the chamber were tracked. Scale bar: 50 µm.
